# Reorganizing the RNA polymerase II complex for replication of an infectious noncoding RNA in vivo

**DOI:** 10.1101/2025.05.12.653476

**Authors:** Jie Hao, Zhuan Qin, Junfei Ma, Jie Qu, Yunhan Wang, Svetlana Y. Folimonova, Bin Liu, Wenwei Li, Ying Wang

**Author notes:** These authors contributed equally to this work (Jie Hao, Zhuan Qin, Junfei Ma, Jie Qu). Current address: Medpace Reference Laboratories, Cincinnati OH 45227, USA.

## Abstract

DNA-dependent RNA polymerases (DdRPs) recognize not only DNA but also RNA templates. This RNA-dependent RNA polymerase (RdRP) activity is exploited by viroids in plants and human hepatitis delta virus in animals. A major knowledge gap exists regarding the molecular basis conferring this RdRP activity. Here, we provide evidence supporting the reorganization of the 12-subunit Pol II to 7-subunit in vivo for PSTVd transcription. Rpb4, Rpb5, Rpb6, Rpb7, and Rpb9 are not involved in PSTVd transcription *in planta*. A splicing variant of transcription factor IIIA with seven zinc finger domains (TFIIIA-7ZF) aids the remodeled Pol II in transcribing PSTVd. Using AlphaFold3, the structure of the remodeled Pol II with PSTVd RNA and TFIIIA-7ZF was generated. The predicted structure and experimental data both show that the N-terminus of TFIIIA-7ZF binds to the left terminal domain of PSTVd, while the C-terminus interacts with Rpb2. Interestingly, AlphaFold3 also predicts the bending at PSTVd loop 8 in the TFIIIA-7ZF/PSTVd complex. Replacing this loop 8 with a rigid double-stranded conformation impairs the TFIIIA-7ZF/PSTVd interaction. Altogether, our demonstrate an active form of heterogenous organization of the essential Pol II enzyme in vivo and provide structural insights into the organization of Pol II transcription complex on RNA template.

## Introduction

Transcription is a fundamental process that mediates the interplay between genetic information and phenotypes and, thus, is vital for organismal growth and responses to environmental cues (Crick, 1958; Crick, 1970; Levine and Tjian, 2003; Spitz and Furlong, 2012). Transcription is catalyzed by DNA-dependent RNA polymerases (DdRPs) that generally polymerize RNA molecules based on DNA templates (Thomas and Chiang, 2006; Ream et al., 2009; Khatter et al., 2017). Interestingly, it is well-known that some DdRPs can recognize both DNA and RNA templates for transcription (Dezelee et al., 1974). The RNA-dependent RNA polymerase (RdRP) activity of DdRPs regulates gene expression in diverse organisms across kingdoms. For instance, under nutrient-deficient conditions, the noncoding 6S RNA of *Escherichia coli* binds to the bacterial DdRP and prevents it from transcribing DNA templates. In response to nutrient availability, the DdRP transcribes a short RNA using the 6S RNA template that leads to the de-sequestration of the DdRP. This step, in turn, enables binding to DNA promoters and the synthesis of mRNAs (Wassarman and Saecker, 2006). RNA polymerase II (Pol II) in mammalian cells binds to a hairpin within the noncoding B2 RNA and uses the longer strand of the hairpin sequence as a template to extend the short strand by transcription. This extension subsequently destabilizes the B2 RNA, representing a novel posttranscriptional mechanism to regulate RNA stability (Wagner et al., 2013). RNA-based pathogens, such as viroids and the human hepatitis delta virus (HDV), alsorely on this RdRP activity for replication in corresponding hosts (Flores et al., 2016). Notably, both (+) and (-) strands of potato spindle tuber viroid (PSTVd, the type speciesof the *Pospiviroidae* family) are transcribed by Pol II. Despite the importance, however, the underlying regulatory mechanism of this RdRP activity of DdRPs remains elusive.

Taking Pol II on DNA templates as an example, this 12-subunit polymerase relies on the coordination of multiple general transcription factors (i.e., TFIIA, TFIIB, TFIID, TFIIE, TFIIF, and TFIIH) during transcription initiation around DNA promoter regions (Roeder, 1998; Lee and Young, 2000; Lemon and Tjian, 2000; Armache et al., 2003). By and blarge, a minimal set of five general transcription factors (TFIIB, TFIID, TFIIE, TFIIF, and TFIIH) is required for the promoter-driven transcription catalyzed by Pol II (Bushnell et al., 1996; Liu et al., 2013). Transcription elongation, particularly in the cellular environment, requires additional factors, including TFIIS, TFIIF, SPT6, etc (Schweikhard et al., 2014; Antosz et al., 2017; Vos et al., 2018b). By contrast, the organization of the transcription complex on RNA templates were poorly defined. Interestingly, emerging evidence implies that Pol II complex on RNA templates may be organized in a distinct fashion.

In a protoplast cell-based replication system, we found that a TFIIS loss-of-function mutant did not impair the propagation of potato spindle tuber viroid (PSTVd) (Dissanayaka Mudiyanselage and Wang, 2020), which replicates in the nucleoplasm and relies on Pol II for de novo transcription (Rackwitz et al., 1981; Wang et al., 2016; Dissanayaka Mudiyanselage and Wang, 2020; Dissanayaka Mudiyanselage et al., 2022; Ma et al., 2024). This observation inspires us to reason that certain general transcription factors may not be involved in the RdRP activity of Pol II. On the other hand, an alternative splicing product of the transcription factor IIIA with seven zinc finger domains (TFIIIA-7ZF) is essential to engage the remodeled Pol II to PSTVd RNA templates for transcription (Wang et al., 2016; Dissanayaka Mudiyanselage and Wang, 2020; Dissanayaka Mudiyanselage et al., 2022). Previous studies showed that TFIIIA-7ZF interacts with Pol II as well as with (+) and (-) PSTVd RNAs in vivo. Overexpression of TFIIIA-7ZF led to the elevated PSTVd titer, while silencing of *TFIIIA-7ZF* reduced PSTVd accumulation (Wang et al., 2016). Moreover, TFIIIA-7ZF is required in a Pol II-based in vitro transcription assay for generating full-length PSTVd RNA products (Wang et al., 2016; Dissanayaka Mudiyanselage and Wang, 2020; Dissanayaka Mudiyanselage et al., 2022). This is akin to the replication of HDV that requires a virus-encoded HDAg-S protein to ensure robust processivity in generating long RNA products (Yamaguchi et al., 2001).

More recently, we found that a “remodeled” Pol II, which is composed of Rpb1, Rpb2, Rpb3, Rpb8, Rpb10, Rpb11, and possibly Rpb12, actively catalyzes the RNA polymerization using PSTVd RNA templates in the Pol II-based in vitro transcription system (Dissanayaka Mudiyanselage et al., 2022). The heterogeneous organization of Pol II echoes some related experimental evidence. First of all, the absence of Rpb9 in the “remodeled” Pol II explains the higher mutation rates of PSTVd during replication (Lopez-Carrasco et al., 2017), when compared to the Pol II fidelity on DNA templates (Gout et al., 2017). Furthermore, distinct Pol II subunits appear to control the transcription of different sets of genes according to degron-based transiently degradation of each subunit in mammalian cell lines (Li et al., 2022). Taken together, evidence supports a model of a distinctive organization of the transcription complex on viroid RNA templates. In spite of the supporting evidence, whether such remodeled Pol II exists in vivo and functions in any natural process remains to be determined. Moreover, the molecular basis remains elusive regarding how TFIIIA-7ZF interacts with the remodeled Pol II for RNA-templated transcription.

To gain insights into the organization of the Pol II transcription complex on RNA template, we tested whether Rpb4, Rpb5, Rpb6, Rpb7, Rpb9 and Rpb12 are involved in binding PSTVd RNA templates in vivo. These subunits, except for Rpb12 that could not be detected due to the small size, were implied to be absent in PSTVd transcription based on in vitro data (Dissanayaka Mudiyanselage et al., 2022). Interestingly, we found that Rpb12 is involved in RNA template binding, while the rest five subunits (Rpb4, Rpb5, Rpb6, Rpb7, and Rpb9) do not interact with PSTVd RNA template in vivo. This observation echoes our recent nLC-MS/MS analysis of the remodeled Pol II on RNA templates, providing compelling evidence supporting the re-organization of Pol II for transcribing viroid RNA templates in plants.

Based on this information, we predicted the structure of the remodeled Pol II with seven subunits (Rpb1, Rpb2, Rpb3, Rpb8, Rpb10, Rpb11, and Rpb12) as well as TFIIIA-7ZF on the (+) PSTVd RNA template. The predicted model provides the opportunity to understand the structural mechanism of an RNA-specific transcription factor (i.e., TFIIIA-7ZF) in RNA-templated transcription. We experimentally tested the model to validate the interaction and structural geometry of RNA templates, TFIIIA-7ZF, and the subunits in the remodeled Pol II. Furthermore, we predicted the complex of TFIIIA-7ZF and (+) PSTVd RNA that pinpoint zinc finger domains (ZF) 2, 3, 6 as possible RNA binding domains. Experimental data clearly support the critical role of ZF3 and ZF6 in RNA binding. Furthermore, the empirical data also showed that the RNA bending at the loop 8 region is critical for TFIIIA-7ZF and PSTVd interaction in the absence of Pol II. Therefore, our analyses pinpoint a structural reorganization of the C-terminal domain of TFIIIA-7ZF from bending the RNA template to bridging the template to polymerase.

Altogether, our data support the in vivo presence of the remodeled Pol II and illustrate how TFIIIA-7ZF bridges the RNA template and the remodeled Pol II, which provides structural insights into the novel regulation of transcription by using an altered organization of polymerase complex.

## Results

### The composition of in vivo Pol II complex on RNA templates

Pol II has been well documented for catalyzing the replication of nuclear-replicating viroids (Wang, 2021; Ma et al., 2023). Our recent findings demonstrate that Pol II can catalyze RNA-templated transcription in vitro without a complete set of 12 subunits (Dissanayaka Mudiyanselage et al., 2022). We termed this distinct organization the “remodeled” Pol II. Those subunits are essential for the cell survival, so we cannot assess the model by using loss-of-function mutants. To corroborate the remodeled Pol II organization in vivo, we performed RNA immunoprecipitation (RIP) analyses to test the interaction between PSTVd RNA and the Pol II subunits absent from the remodeled Pol II (i.e., Rpb4, Rpb5, Rpb6, Rpb7, and Rpb9), using Rpb3 as a positive control. We could not detect Rpb12 in our previous work (Dissanayaka Mudiyanselage et al., 2022), but it was suspected that the small size of Rpb12 rendered it being filtered out by size cut-off columns during sample preparation. Therefore, we also included Rpb12 in this test. Since most of these subunits have multiple gene copies in plants, we selected the gene copies that are expressed and incorporated into Pol II based on published data (Ream et al., 2015). These subunits were expressed in the PSTVd-infected *N. benthamiana* plants as fusions with a GFP-tag for RIP analyses. Plants infiltrated with agrobacteria without the expression vector served as negative control.

As shown in Fig.1A, GFP-trap successfully precipitated all the target subunits. We then used RT-qPCR to detect PSTVd RNA templates enriched in the immunoprecipitated fraction. 5.8S rRNA was used as negative control for normalizing RT-qPCR data. As shown in Fig. 1A, PSTVd RNA was significantly co-precipitated in samples expressing subunits Rpb3 and Rpb12, confirming that Rpb3 and Rpb12 are part of the remodeled Pol II in vivo. By contrast, PSTVd RNA was not significantly enriched in samples expressing subunits Rpb4, Rpb5, Rpb6, Rpb7, or Rpb9 as compared with the negative control samples. Therefore, these subunits are unlikely part of the remodeled Pol II on PSTVd RNA templates. The results are consistent with our previous in vitro analysis, except for Rpb12. As previously speculated, Rpb12 was likely lost during our sample preparation for nLC-MS/MS analysis, which was reported in a similar study (Antosz et al., 2017).

**Figure 1.**
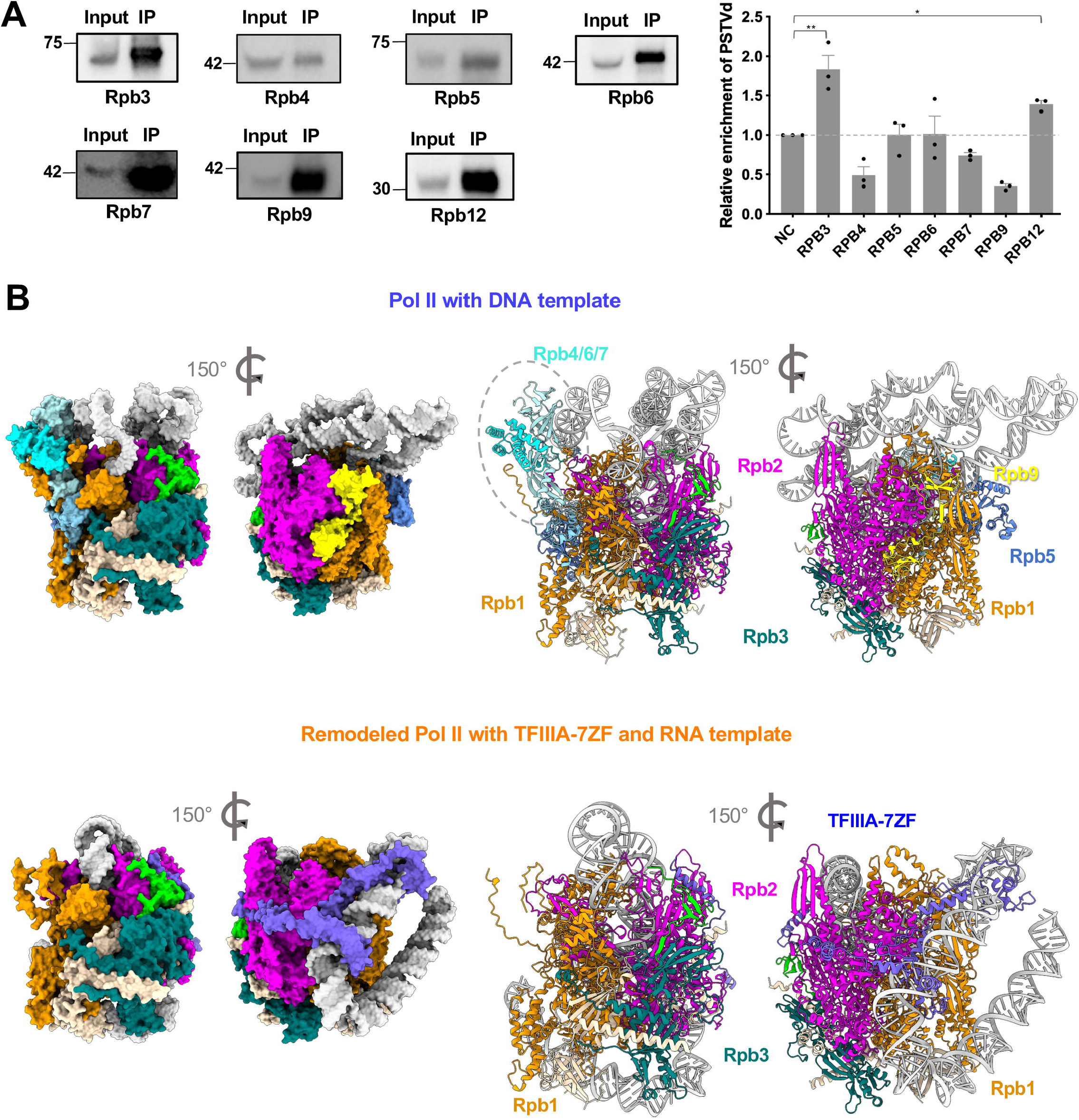
Remodeled Pol II in vivo and its predicted structure. **A)** RNA-immunoprecipitation using Pol II subunits. GFP-tagged Rpb3, Rpb4, Rpb5, Rpb6, Rpb7, Rpb9, and Rpb12 were subject to RNA immunoprecipitation to test their in vivo interaction with PSTVd RNA templates. Numbers indicate the molecular weight (Kda). RNAs from input and immunoprocipitated (IP) fraction were used for RT-qPCR analysis. 5.8S rRNA was used as negative control to normalize PSTVd enrichment. Data from there replicates were used for T-test analysis. **B)** Predicted structures of Pol II with DNA template and remodeled Pol II with TFIIIA-7ZF and RNA template. The catalytic subunits (Rpb1 and Rpb2) as well as the distinct subunits (Rpb4/5/6/7/9) and TFIIIA-7ZF are color coded. The stalk region is highlighted in a dash-line circle.

### Structure overview of the TFIIIA-7ZF/Pol II complex on the (+) PSTVd RNA templates

Previous structural studies have resolved multiple high-quality Pol II complexes with general transcription factors at different transcription stages (Cramer et al., 2001; Kettenberger et al., 2004; Plaschka et al., 2015; Robinson et al., 2016; Ehara et al., 2017; Vos et al., 2018b; Vos et al., 2018a; Dienemann et al., 2019). The structure of a canonical TFIIIA protein with nine zinc finger domains (ZFs) is also available (PDB 1tf6) (Nolte et al., 1998). However, the structure illustrating the complex of Pol II and RNA-specific transcription factors (i.e., HDAg-S and TFIIIA-7ZF) is lacking. Nevertheless, these resolved structures provided the basis for predicting a Pol II complex with other factors. Taking the advantage of the emergence of AlphaFold3 (Abramson et al., 2024), we predicted the structure of the TFIIIA-7ZF/Pol II complex on the (+) PSTVd RNA template (359 nt)(Movie 1). For comparison, we also predicted the Pol II complex (with 12 subunits) on a similar length of a DNA template (395 bp), using the sequence from an expression vector pRTL2 with the 35S promoter. As shown in Fig. S1, the predicted models have very high confidence for the Pol II and remodeled Pol II complexes, which reflects the presence of high-quality Pol II data. The structures for the nucleic acid portions generally have low confidence, which is expected since AlphaFold3 is generally thought to have less accuracy in predicting nucleic acid structures.

The remodeled Pol II lacks the “Stalk” that consists of Rpb4/7 and its adjacent Rpb6 (Fig. 1B). This absence is possibly not directly caused by TFIIIA-7ZF occupancy because TFIIIA-7ZF does not share any common binding sites with these three subunits (Fig. 1B). The absence of Rpb5 and Rpb9 may be caused by TFIIIA-7ZF occupancy since their binding sites crossover with the TFIIIA-7ZF binding site on the surface of Rpb1.

### Testing the geometry of TFIIIA-7ZF in the Pol II/TFIIIA-7ZF/PSTVd complex

We noticed that TFIIIA-7ZF wraps the RNA template while attaches to the remodeled Pol II complex (Fig. 1B). The N-terminal region of TFIIIA-7ZF is involved in binding with the RNA template, and the TFIIIA-7ZF C-terminus appears to be critical for binding with the Pol II core, particularly Rpb2. As shown in Fig. 2A, residues R243, R245, and K257 of TFIIIA-7ZF are predicted to be involved in binding with Rpb2 Q91 in the Protrusion domain and E549/E550 in the External 2 domain. Residues R243, R245, and K257 are outside of the seven zinc finger domains towards the C-terminus of the TFIIIA-7ZF. We also noticed that the C-terminus end of TFIIIA-7ZF is relatively close to the N-terminus of Rpb12 but away from the N-terminus of Rpb3 (Fig. 2B).

**Figure 2.**
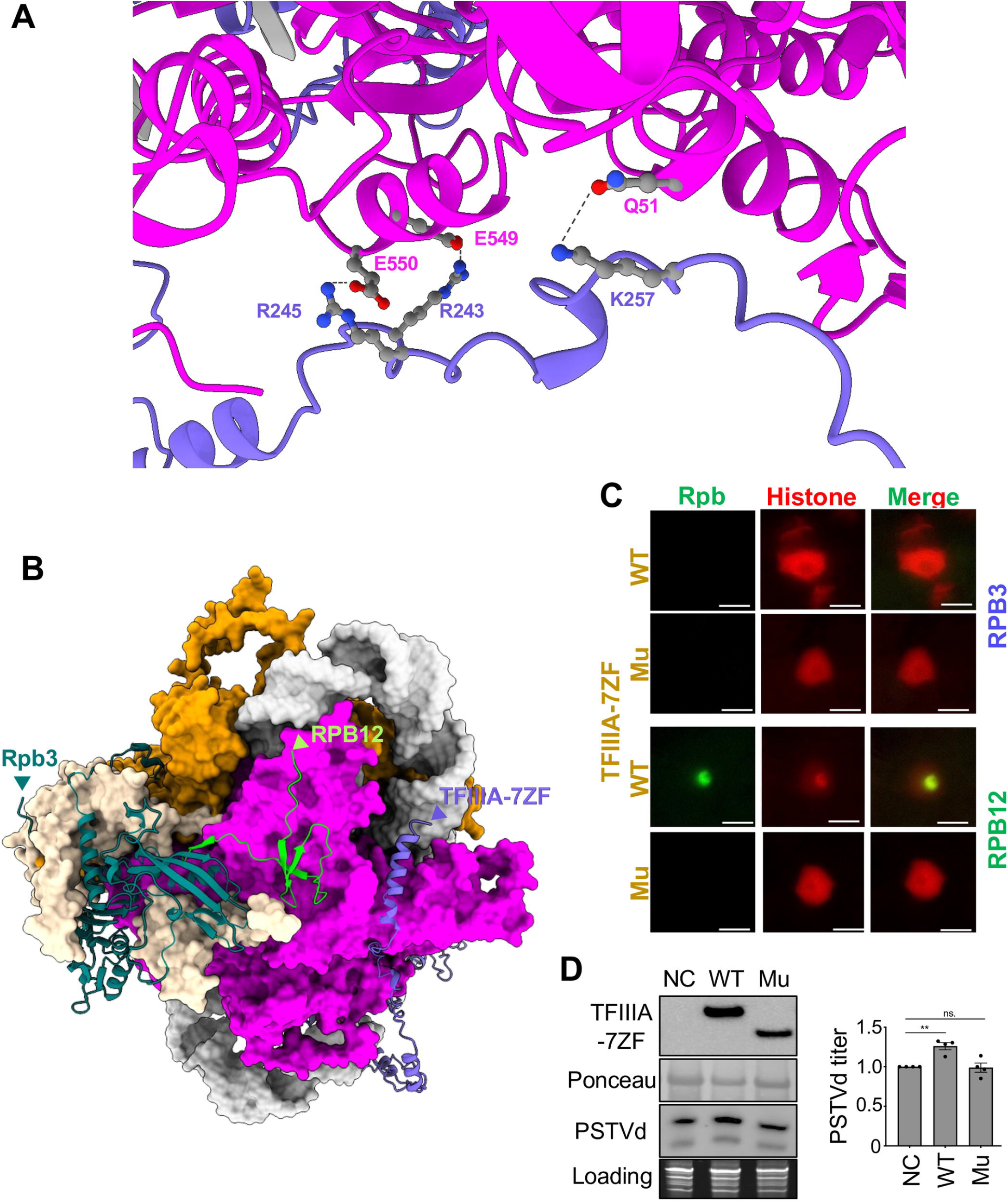
Relative position of TFIIIA-7ZF and Pol II subunits. **A)** Predicted interactions between Rpb2 and TFIIIA-7ZF. Rpb2 and TFIIIA-7ZF are depicted in magenta and blue, respectively. B) Predicted structure showing the relative position between the C-terminus of TFIIIA-7ZF (labeled by an arrow head) and the N-terminus of Rpb3 and Rpb12 (labeled by arrow heads). **C)** Bi-fluorescence complementation (BiFC) confirms that the C-terminus of TFIIIA-7ZF is close to the N-terminus of Rpb12 but not the N-terminus of Rpb3. Mu, TFIIIA-7ZF^C-del^. Scale Bar, 10 μm. **D)** RNA gel blots showed that transiently over-express WT TFIIIA-7ZF, but not TFIIIA-7ZF^C-del^, enhanced PSTVd accumulation in planta. TFIIIA-7ZF proteins were detected by α-HA antibody. Ponceau S staining serves as loading control for immunoblots. PSTVd was detected by RNA gel blots, and ethidium bromide staining of rRNAs serves as the loading control.

To empirically test the predicted structural geometry of the TFIIIA-7ZF/Pol II complex, we used bimolecular fluorescence complementation assay (BiFC) to assess the relative distance between TFIIIA-7ZF (with a C-terminal half of YFP fused to the end of the TFIIIA-7ZF C-terminus) and Rpb12 or Rpb3 (with a N-terminal half of YFP fused to the subunit’s N-terminus). As shown in Fig. 2C, only the Rpb12/TFIIIA-7ZF pair, but not the Rpb3/TFIIIA-7ZF pair, could complement the YFP fluorescence. We then designed a C-terminus-lacking variant of TFIIIA-7ZF (TFIIIA-7ZF^C-del^; aa 1-228) that still keeps all seven zinc finger domains but lacks the critical residues for Rpb2 binding. This TFIIIA-7ZF^C-del^ could not complement YFP fluorescence with Rpb12 (Fig. 2C). Furthermore, TFIIIA-7ZF^C-del^ could not enhance PSTVd accumulation as compared with WT TFIIIA-7ZF (Fig. 2D). These results confirm that the C-terminus of TFIIIA-7ZF is functionally important for binding with the remodeled Pol II complex. Importantly, the BiFC data corroborate the predicted structure of the complex in terms of the relative distance between TFIIIA-7ZF and Pol II subunits (i.e., Rpb3 and Rpb12).

### Structure of TFIIIA-7ZF/viroid complex

In our Pol II-based in vitro transcription assay, incubating RNA template with TFIIIA-7ZF prior to the addition of Pol II is critical for optimized transcription efficiency (Dissanayaka Mudiyanselage and Wang, 2020), which implies that TFIIIA-7ZF and PSTVd RNA-protein complex is formed first before recruiting Pol II. However, it is unclear how TFIIIA-7ZF coordinates its seven ZFs for RNA template binding. Therefore, we were interested in understanding the structure of TFIIIA-7ZF and the PSTVd (+) genomic RNA by using AlphaFold3. Previously, it was speculated that the left terminal region of PSTVd (+) RNA genome forms either a rod-shape structure or a bifurcated structure during transcription (Dissanayaka Mudiyanselage et al., 2018). However, the model predicted by AlphaFold3 provided an unexpected conformation (Fig. 3A). While PSTVd RNA bases still pair similarly as in the rod-shape structure, the TFIIIA-7ZF protein bends the overall rod-shape structure that leads to the formation of an enlarged left terminal region. We noticed that base A51, which resides in loop 8, is right at the bending site. Using an established mutant that changes loop 8 to a double-stranded conformation (Zhong et al., 2008), we demonstrated that the rigid double-stranded conformation impairs PSTVd RNA binding with TFIIIA-7ZF (Figure 3C), which is in support of bending at loop 8 in the TFIIIA-7ZF/PSTVd complex. It also aligns with the previous observation that this loop 8 mutant exhibited reduced replication efficiency (Zhong et al., 2008). Together, the data support the importance of RNA bending for interacting with TFIIIA-7ZF protein.

**Figure 3.**
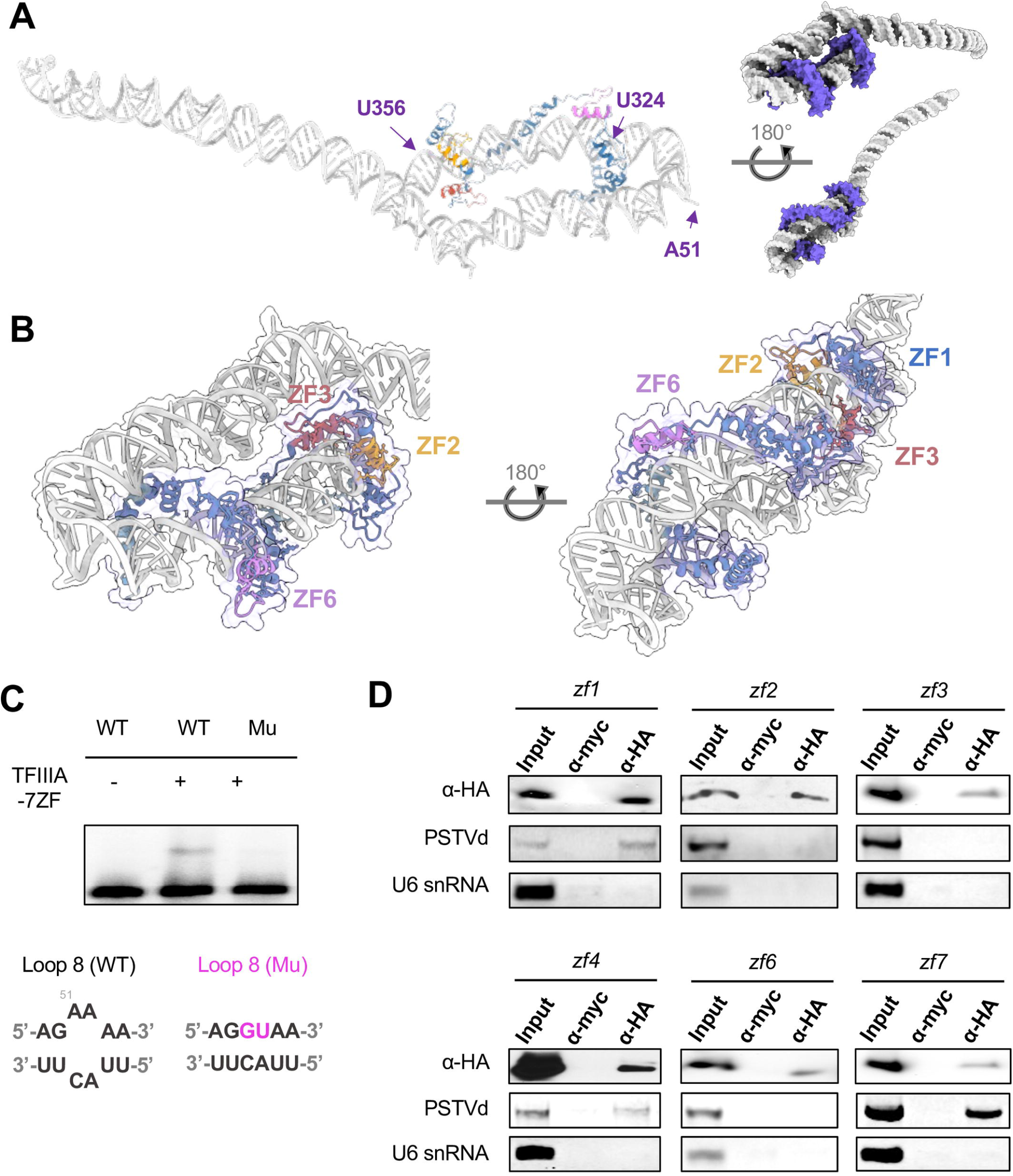
Structure of TFIIIA-7ZF and (+) PSTVd RNA. **A)** Predicted structure of TFIIIA-7ZF and PSTVd RNA. Different zinc finger domains are color highlighted. Regions between U356 and U324 are predicted to be covered by TFIIIA-7ZF. **B)** A zoom-in view of the critical zinc fingers for RNA binding. **C)** EMSA demonstrating that loop 8, which is predicted as the bending site, is critical for TFIIIA-7ZF binding. **D)** RNA-immunoprecipitation demonstrating the function of the zinc fingers. HA-tagged proteins were expressed in PSTVd-infected plants. α-HA and α-MYC were used for immunoprecipitation (IP) and control treatments, respectively. Semi quantitative RT-PCR were used for examining PSTVd enrichment in IP fractions, with U6 snRNA served as negative control. All experiments were repeated at least twice.

A close look at the predicted structure showed that ZF2 and ZF3 clamp the major groove of the RNA template (Fig. 3B)., The C-terminus of TFIIIA-7ZF binds to the RNA template, instead of binding with Rpb2 in the Pol II/TFIIIA-7ZF/PSTVd complex shown in Fig. 2A. The TFIIIA-7ZF binding sites are estimated as G328 to G354 (Fig. 3A), akin to our foot-printing assay that mapped the binding region from U331 to G346 (Wang et al., 2016). This region is also proximal to the transcription initiation site between U359 and C1 (Kolonko et al., 2006). The significant overlapping between the experimental data and the prediction model suggests that the predicted model can indeed reflect the structure of the (+) PSTVd/TFIIIA-7ZF complex. To further map the ZFs that are critical for RNA templates binding, we introduced a point mutation changing the first histidine to asparagine to disrupt each C2H2 ZF as previously reported (Lee et al., 2006; Rothfels et al., 2007) and performed RIP using these variants with an HA tag. We could not detect the expression of the *zf5* variant in plants, potentially attributable to poor stability of the protein. Nevertheless, the rest six variants were successfully expressed and detected in the eluted RIP fractions. Since our primer sets can detect both (+) and (-) PSTVd strands simultaneously in one-step RT-qPCR reactions, we only used semi-quantitative RT-PCR to test for the enrichment of the (+) PSTVd RNA in the immunoprecipitated fractions. As shown in Fig. 3D, the *zf2, zf3*, and *zf6* variants consistently failed to precipitate PSTVd RNA, indicating that these domains are critical for RNA binding, which provides a strong support for the predicted model of the RNP.

To further test the critical zinc fingers involved in (+) PSTVd RNA binding, we performed electrophoretic mobility shift assays (EMSAs) using each *zf* variant. As shown in Fig. 4, disruption of either ZF3 or ZF6 significantly impacts RNA binding, as these variants lost the ability to achieve a complete RNA shifting in comparison to wildtype and other *zf* variants. Interestingly, the rest of *zf* variants demonstrated similar or even higher affinity to the PSTVd RNA as indicated by the dissociation constant (Kd). Notably, the *zf2* variant could bind with (+) PSTVd RNA in EMSA tests, which is different from the RIP outcomes presented in Fig. 3D. This discrepancy implies that the ZF2 domain may be involved in other interactions in the cellular environment. On the other hand, when ZF2 is impaired, ZF1 might contribute to stabilizing RNA binding in vitro through working together with ZF3 for RNA binding. This is implied by the geometry between ZF1 and ZF3 on PSTVd RNA in the predicted model (Fig. 3B). In spite of this discrepancy of ZF2, RIP, EMSA, and the structural model all support the critical roles of ZF3 and ZF6 in (+) PSTVd binding.

**Figure 4.**
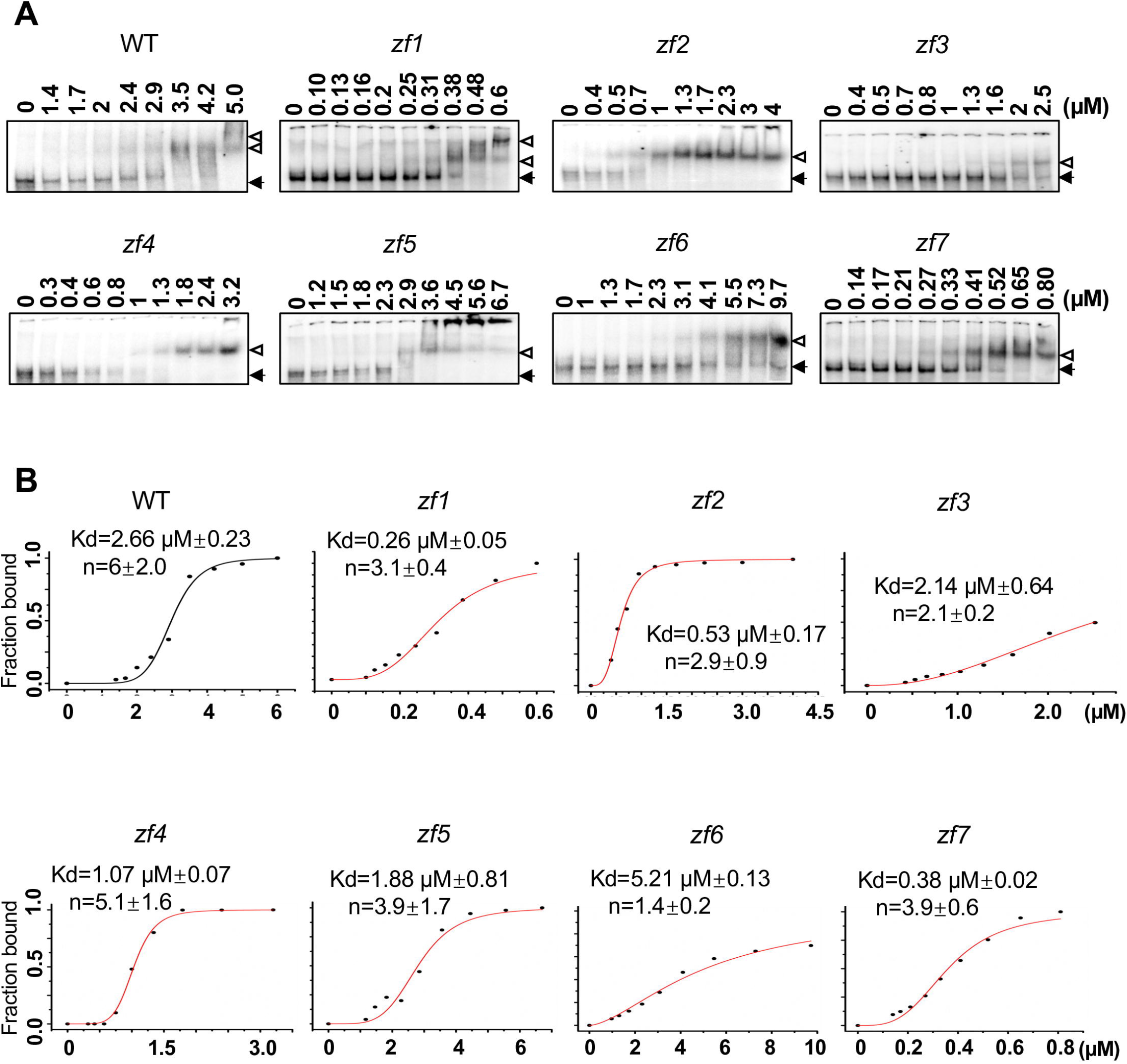
Interaction of TFIIIA-7ZF variants with PSTVd in EMSAs. **A)** Representative EMSA result for the binding between TFIIIA-7ZF zinc finger variants and PSTVd. Protein concentrations are listed for each lane. **B)** Representative binding curves for each TFIIIA-7ZF variants. Each data point was calculated based on three replicates. Protein concentrations are listed on X axes.

## Discussion

The results presented in this study demonstrate a distinct organization of Pol II transcribing viroid RNA template in planta, a new layer of transcriptional regulation in a natural biological process. By using RNA-immunoprecipitation, we showed that only a subset of Pol II subunits are associated with PSTVd RNA, in line with of our previous results from an in vitro setting (Dissanayaka Mudiyanselage et al., 2022). Rpb1, Rpb2, Rpb3, Rpb8, Rpb10, Rpb11, and Rpb12 constitute the remodeled Pol II for RNA-templated transcription. During the assembly of Pol II complex, there are several intermediate complexes/subassemblies, including the Rpb1 subassembly containing Rpb1/5/6/8, the Rpb2 subassembly containing Rpb2/9, the Rpb3 subassembly containing Rpb3/10/11/12 and the Rpb4/7 subassembly (Stalk)(Wild and Cramer, 2012). Interestingly, the remodeled Pol II contains the intact Rpb3 subassembly, a modified Rpb1 subassembly lacking Rpb5 and Rpb6, and a modified Rpb2 subassembly lacking Rpb9 (Boulon et al., 2010; Gomez-Navarro et al., 2013). The stalk portion of the Pol II is completely missing in the remodeled Pol II. It is known that Rpb4, Rpb5, Rpb7, and Rpb9 tend to dissociate from the Pol II complex upon alpha-Amanitin treatment (Boulon et al., 2010), suggesting that these subunits are prone to leave the Pol II complex under stress. Our findings also align with a recent finding of possible heterogeneous organization of Pol II in mammalian cells (Li et al., 2022), suggesting that the transcriptional regulation at the polymerase organization level is more complex than previously recognized.

Confirming the remodeled Pol II organization in vivo paves the way for further investigations on the different regulations of DNA-dependent versus RNA-templated transcription. For instance, the Pol II stalk is involved in binding with capping enzymes for adding G-cap to nascent mRNAs (Garg et al., 2023; Li et al., 2024). This explains the lack of G-cap in viroid RNAs (Ma et al., 2022a). Whether HDV RNA-templated transcription also uses a remodeled Pol II is not immediately clear. Some HDV-derived mRNAs contain a G-cap, but Pol II-transcribed genomic or antigenomic RNAs do not (Gudima et al., 2000; Taylor, 2015).

Although the detailed structural basis has largely been determined for almost every step in RNA polymerases transcribing DNA templates, the molecular basis underlying the widely existing RNA-templated transcription remains elusive. To date, the only known structure, which contained 12-subunit Pol II, TFIIS, and a synthetic mosaic RNA template with partial HDV sequence, was determined by using a crystallographic analysis (Lehmann et al., 2007). This structure did not include the RNA-specific transcription factor (HDAg-S) for HDV RNA templates. In addition, this complex does not possess strong processivity for yielding long products (longer than 100 nt) that occur in nature. Moreover, the TFIIS-based cleavage allows Pol II to catalyze primed transcription on some RNA templates, but it is not required for transcribing PSTVd as Pol II catalyzes de novo transcription using PSTVd RNA template (Dissanayaka Mudiyanselage and Wang, 2020). Therefore, alternative structure model is needed for understanding RNA-templated transcription.

Here, we obtained predicted structures of the transcription complex and experimentally tested the structural model. In the predicted TFIIIA-7ZF/PSTVd structural model, notably, TFIIIA-7ZF binds around the left terminal part of PSTVd (+) RNA template and bends the RNA to form an enlarged tertiary structure (Fig. 3). The loop 8 mutant that makes the bending region more rigid impairs TFIIIA-7ZF and PSTVd interaction, supporting the functional relevance of this RNA bending. This enlarged overall structure possibly contributes to the recruiting of Pol II. The predicted model also suggests that the viroid RNA promoters may not entirely rely on primary sequences. Instead, an overall bended tertiary structure is likely critical for transcription. This organization is different from previously reported hairpin-like RNA promoter used for testing a chimeric HDV RNA template (Lehmann et al., 2007). This model may help to explain that over 30 nuclear-replicating viroids with distinct sequences all use Pol II for replication (Wang, 2021). When Pol II is recruited to the PSTVd/TFIIIA-7ZF complex, the C-terminus of TFIIIA-7ZF somehow moves from RNA binding to the association with Rpb2. A C-terminus deletion impairs TFIIIA-7ZF binding with the remodeled Pol II and abolishes its function in aiding PSTVd accumulation. Therefore, these data outline a critical structural transition of the TFIIIA-7ZF C-terminus from bending the RNA template to connecting the RNA template with the polymerase. The details of the structural transition, from the association with the bended RNA template to the Rpb2 in the remodeled Pol II, deserve future investigations.

In the Pol II-based in vitro transcription assay, factors that are needed for DNA-dependent transcription elongation in vivo (i.e., TFIIF, RTF1, PAF1, LEO1, CTR9, SPT6, SPT16, and SSRP1-B), were all absent in the remodeled Pol II on PSTVd RNA template (Dissanayaka Mudiyanselage et al., 2022). Instead, a specialized RNA-specific transcription factor, TFIIIA-7ZF, enhances Pol II processivity on PSTVd RNA templates (Wang et al., 2016; Dissanayaka Mudiyanselage and Wang, 2020; Dissanayaka Mudiyanselage et al., 2022). In the simplest experimental setup, products with hundreds of nucleotides were generated in vitro by simply mixing Pol II purified from plants, purified TFIIIA-7ZF expressed in bacteria, and synthesized PSTVd RNA templates (Wang et al., 2016; Dissanayaka Mudiyanselage and Wang, 2020; Dissanayaka Mudiyanselage et al., 2022). However, whether any general transcription factors (in addition to TFIIIA-7ZF) or cofactors are needed for PSTVd RNA-templated transcription in cells remains to be determined. We also tested TFIIF and TFIIS in our RIP analyses. As shown in Fig. S2, there was no evidence of any enrichment of PSTVd in the TFIIS immunoprecipitated fraction, which is consistent with our previous data that TFIIS is dispensable for PSTVd replication (Dissanayaka Mudiyanselage and Wang, 2020). Interestingly, we observed significant enrichment of PSTVd in TFIIF-immunoprecipitated fraction (Fig. S2), suggesting that TFIIF could be involved in RNA-templated transcription. It remains to be determined whether TFIIF is involved in the initiation or elongation step in RNA-templated transcription. Nevertheless, this observation also demonstrates that Pol II may work with common and unique sets of general transcription factors when transcribing DNA and RNA templates.

## Materials and Methods

### Plant Materials

*Nicotiana benthamiana* plants were grown in a growth room with 28°C and a 14-h/10-h light/dark cycle. During agroinfiltration period, plants were moved into a growth room with 22°C and a 14-h/10-h light/dark cycle. PSTVd-infection and validation steps were the same as described previously (Zhu et al., 2002; Ma et al., 2022b).

### Molecular Cloning

Following clones were ordered from Arabidopsis Biological Resource Center (ABRC) at the Ohio State University: Rpb4 (DKLAT5G09920) and Rpb7 (DKLAT5G59180). ORFs of Rpb5 (At3g22320), Rpb6 (At2g04630), Rpb9 (At4g16265), Rpb12 (At5g41010),

TFIIF1 (At1g75510), and TFIIS (AT2G38560) were cloned via conventional RT-PCR and inserted into the pENTR-D-TOPO vector (ThermoFisherSci, Waltham, MA) and recombined to destination vectors. All clones were verified by Sanger Sequencing. Primers were listed in the Supp Table 1. Loop 8 mutant was reported previously (Zhong et al., 2008). TFIIIA-7ZF WT and point mutation variants were reported previously (Dissanayaka Mudiyanselage et al., 2022). The C-terminal deleted TFIIIA-7ZF mutant (aa 1-228) was cloned into the pENTR-D-TOPO vector using WT clone as template, followed by LR recombination to festination vectors.

### RNA-Immunoprecipitation (RIP)

RIP was performed following our established protocol (Wang et al., 2016). PSTVd-infected *N. benthamiana* plants were agro-infiltrated with destination vectors to express GFP-tagged Pol II subunits, TFIIF, and TFIIS, as well as HA-tagged TFIIIA-7ZF variants. Three days post infiltration, protein expressions were verified by microscopy observation of green fluorescence before RIP procedures. Collected leaf samples were lysed in RIP buffer (25 mM Tris pH 7.5, 150 mM NaCl, 0.5 mM EDTA, 10% glycerol, 0.1% Triton X-100, 0.2% NP-40, protease inhibitor cocktail following manual) as described before (Wang et al., 2016). The lysates were incubated with GFP-trap beads (ProteinTech, Rosemont, IL) anti-HA magnetic beads (ThermoFisherSci) or anti-Myc magnetic beads (ThermoFisherSci) for 2 hr at 4°C with rotation. The beads were washed twice in IP buffer and once in RNase-Free water, before elution using acidic Glycine buffer following manual instructions (ThermoFisherSci). The protein fractions were analyzed via immunoblotting as described before in details (Wang et al., 2016). The precipitated RNAs were analyzed by semi-quantitative RT-PCR or RT-qPCR (please see Supp Table 1 for primer sequences). For RT-qPCR, 5.8S rRNA was used as negative control for normalizing PSTVd enrichment folds. Data from three biological replicates were used for T-test analysis.

### Bimolecular fluorescence complementation (BiFC) and Microscopy

For BiFC, PSTVd-infected *N. benthamiana* seedlings were used for agroinfiltration of various combinations of constructs, all including 35S:RFP-Histone 2B (Ma et al., 2022b) as the nucleus marker. The N split (aa 1–174) YFP was fused in front of the N-terminus of RPB3 or RPB12. The C split YFP (aa 175-end) was fused after the C-terminus of TFIIIA-7ZF and its variant. We analyzed 10 randomly chosen regions of infiltrated leaves from at least three plants for each treatment. ECHO Revolve microscope (Discover Echo Inc., San Diego, CA) was used for observing the fluorescence signals.

### Electrophoretic mobility shift assays (EMSA)

For EMSA, TFIIIA-7ZF and its variants were expressed using IMPACT system (New England Biolabs, Ipswich, MA) in bacteria and purified using Chitin beads as previously described (Wang et al., 2016). Protein-RNA binding reactions (20 mM Tris-HCl pH 7.5, 35 mM KCl, 5 µM ZnCl_2_, 3.5 mM MgCl_2_, 10 nM yeast tRNA and 10% (v/v) glycerol) were performed at 25°C using ^32^P-labeled RNAs (20,000 dpm) with increasing concentrations of recombinant proteins (described in Fig. 4). Electrophoresis was carried out on ice using 6% (w/v) polyacrylamide (29:1) gels for 2 h at 120 V with Tris borate buffer (65 mM Tris, 22.5 mM boric acid, pH 8). PhosphorImager cassette was used to visualize the signal in a Personal Molecular Imager (Bio-Rad Laboratories, Hercules, CA). The intensity of each band on the gel was quantified by using Quantity One software (Bio-Rad Laboratories). The binding curves were obtained by plotting the fraction of RNA bound with proteins as previously described (Wang et al., 2016).

## Acknowledgments

We are grateful for the technical support of Shachinthaka D. Dissanayaka Mudiyanselage. This work was supported by US National Science Foundation MCB-2350392 and IOS-2410009 (Ying W), as well as US National Institutes of Health R01AI194920-01 (WL). Yunhan Wang is partially supported by CALS Deans’ scholarship from University of Florida.

## Data and materials availability

All data are available in the main text or the supplementary materials.

---

**Supplementary Table 1. Primer sequences.**

**Supplementary Figure 1. AlphaFold 3 confident scores.**

**Supplementary Figure 2. RNA-immunoprecipitation using TFIIF and TFIIS.**

**Movie 1. Transcription complex containing remodeled Pol II, TFIIIA-7ZF, and (+) PSTVd RNA.**

**Supplementary Dataset 1. AlphaFold3 predicted transcription complex containing remodeled Pol II, TFIIIA-7ZF, and (+) PSTVd RNA.**

**Supplementary Dataset 2. AlphaFold3 predicted transcription complex containing Pol II and a DNA fragment.**

**Supplementary Dataset 3. AlphaFold3 predicted complex containing TFIIIA-7ZF and (+) PSTVd RNA.**

