## Supplementary material for "Reorganizing the RNA polymerase II complex for replication of an infectious noncoding RNA in vivo": Fig. S1

### Pol II with DNA template

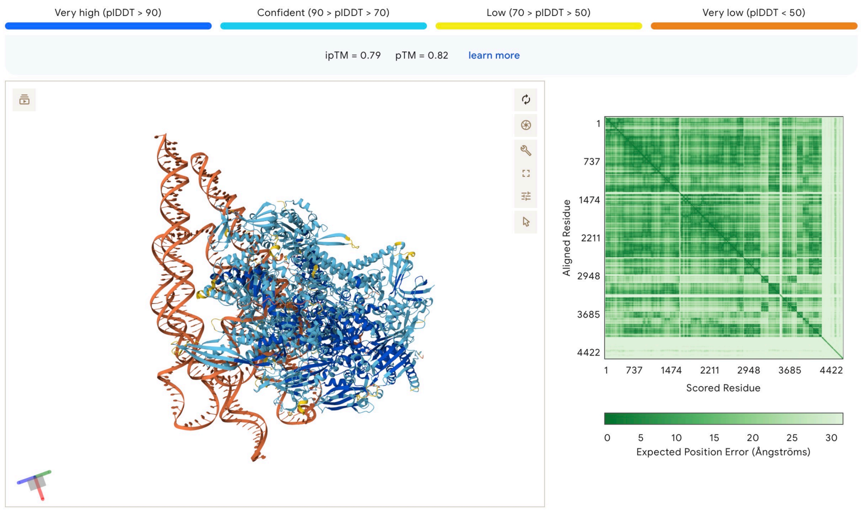


### Remodeled Pol II with PSTVd and TFIIIA-7ZF

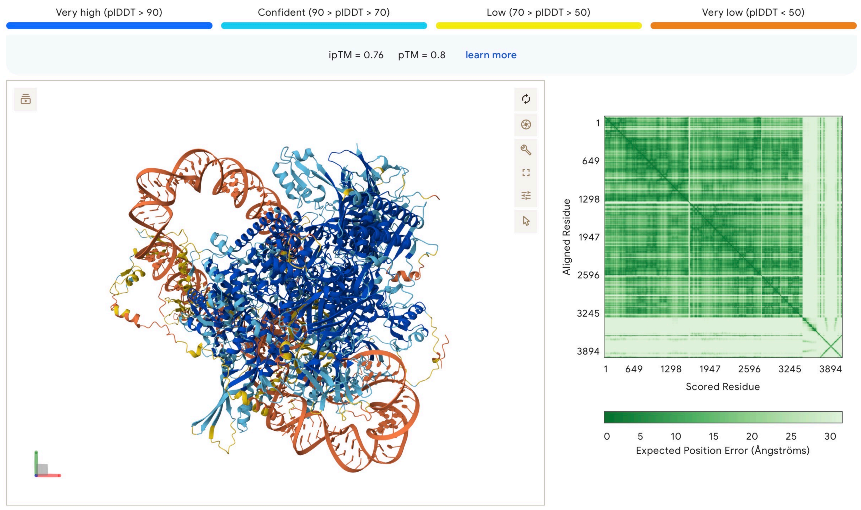


### TFIIIA-7ZF and PSTVd

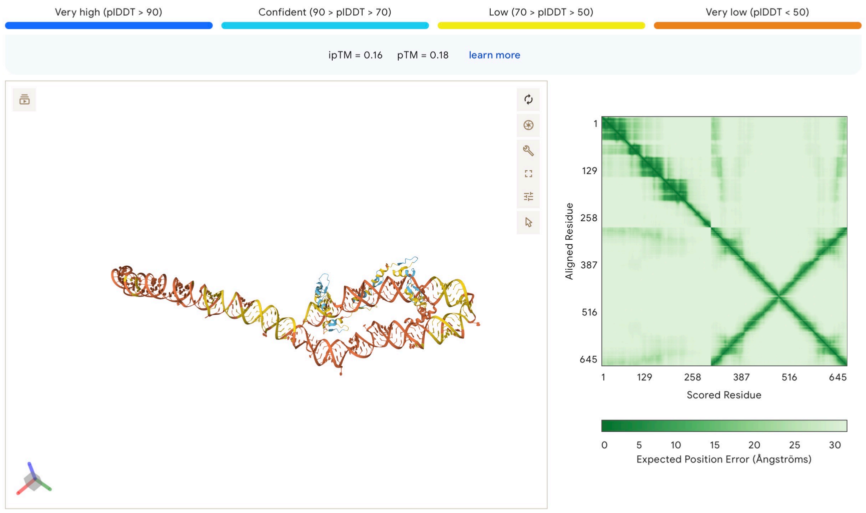


**Figure S1.** AlphaFold 3 confident scores
