## Supplementary material for "Reorganizing the RNA polymerase II complex for replication of an infectious noncoding RNA in vivo": Fig. S2

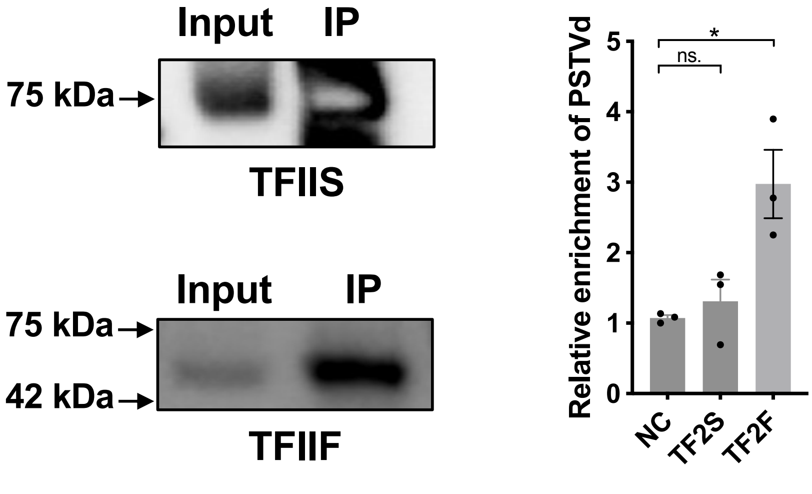


**Figure S2. RNA-immunoprecipitation using TFIIF and TFIIS.** GFP-tagged proteins were subject to RNA immunoprecipitation to test their in vivo interaction with PSTVd RNA templates. RNAs from input and immunoprocipitated (IP) fraction were used for RT-qPCR analysis. 5.8S rRNA was used as negative control to normalize PSTVd enrichment. Data from three replicates were used for T-test analysis.
