## Supplementary material for "Reorganizing the RNA polymerase II complex for replication of an infectious noncoding RNA in vivo": Table S1

Supplementary Table1. Primers

| Primer name | | Sequence |
|---|---|---|
| RPB5 | F | caccATGTTGACGGAAGAGGAGCT |
| | R | TTATACAACATAACGATAGGTAACATAAC |
| RPB6 | F | caccATGGCTGACGACGATTACAATGAAG |
| | R | TCAATCACCACCGACTTGACGTTTC |
| RPB9 | F | caccATGTTTTCTATTGGATTAAAGCTCCAACTT |
| | R | CTATTCTCTCCAACGGTGACTA |
| RPB12 | F | caccATGGATCCAGCGCCCGAACC |
| | R | TCAGCGAGCTTCGTATTGAACAACT |
| TFIIF1 | F | caccATGGAAGATATTCATAATCTCGATATAGAG |
| | R | CTGCCCACCTGTATCATCTTCAG |
| TFIIS | F | caccATGGAGAGTGATTTGATTGATTTGTTCG |
| | R | TCAACAGAACTTCCAGTGGTTGTCAC |
| PSTVd | F | ggggaaacctggagcgaactgg |
| | R | cccggggatccctgaagcgctcc |
| 5.8S rRNA | F | AAACGACTCTCGGCAACG |
| | R | GCGTGACGCCCAGGCAGA |
| U6 snRNA | F | gtcccttcggggacatccgata |
| | R | tttggaccatttctcgatttgtgcgtg |
| TFIIIA-7ZF | zf1 F | CACTTGACGAGAaATCTCTTGCAGC |
| | zf1 R | GCTGCAAGAGATTTCTCGTCAAGTG |
| | zf2 F | CAACATGACTCGGaATGTCAATGAGATGC |
| | zf2 R | GCATCTCATTGACATtCCGAGTCATGTTG |
| | zf3 F | GCATCCAAATTAAAGAAAaATGAGGATTCTC |
| | zf3 R | GAGAATCCTCATTTTTCTTTAATTTGGATGC |
| | zf4 F | TGCCTCAAGGAAaACGTGGAGAGTTG |
| | zf4 R | CAACTCTCCACGTtTTCCTTGAGGCA |
| | zf5 F | GAATATTAAGCGGaATCTCCGTACGCATG |
| | zf5 R | CATGCGTACGGAGATTCCGCTTAATATTC |
| | zf6 F | ATCAAATCTTATTCAGaACGTCAAAGCTG |
| | zf6 R | CAGCTTTGACGTtCTGAATAAGATTTGAT |
| | zf7 F | CGTGAGAGATAGAaATGAAAAGTCTGGC |
| | zf7 R | GCCAGACTTTTCATTTCTATCTCTCACG |
| | WT F | caccATGCAAGAGAGGCCATTTGCATGC |
| | C-del R | GAAATCACCAGGAGTATAAACATG |
